# Lateral and longitudinal compaction of PRC1 overlap zones drive stabilization of interzonal microtubules

**DOI:** 10.1101/2023.01.30.526324

**Authors:** Carline Fermino do Rosário, Ying Zhang, Jennifer Stadnicki, Jennifer L. Ross, Patricia Wadsworth

## Abstract

During anaphase, antiparallel overlapping midzone microtubules elongate and form bundles, contributing to chromosome segregation and the location of contractile ring formation. Midzone microtubules are dynamic in early but not late anaphase; however, the kinetics and mechanisms of stabilization are incompletely understood. Using photoactivation of cells expressing PA-EGF-α-tubulin we find that immediately after anaphase onset, a single highly dynamic population of midzone microtubules is present; as anaphase progresses, both dynamic and stable populations of midzone microtubules coexist. By midcytokinesis, only static, non-dynamic microtubules are detected. The velocity of microtubule sliding also decreases as anaphase progresses, becoming undetectable by late anaphase. Following depletion of PRC1, midzone microtubules remain highly dynamic in anaphase and fail to form static arrays in telophase despite furrowing. Cells depleted of Kif4a contain elongated zones of PRC1 and fail to form static arrays in telophase. Cells blocked in cytokinesis form short PRC1 overlap zones that do not coalesce laterally; these cells also fail to form static arrays in telophase. Together, our results demonstrate that dynamic turnover and sliding of midzone microtubules is gradually reduced during anaphase and that the final transition to a static array in telophase requires both lateral and longitudinal compaction of PRC1 containing overlap zones.

## Introduction

Chromosome segregation during mitosis is accomplished by the bipolar mitotic spindle, which is comprised of microtubules and numerous structural and regulatory proteins (for review see Glotzer, 2009; McIntosh, 2017; Vukušić et al., 2019). A defining characteristic of the mitotic spindle is its dynamic nature. As cells enter mitosis, the nucleation and turnover of microtubules both increase (Heald & Khodjakov, 2015; Khodjakov & Rieder, 1999; Rusan et al., 2001) and this dynamic behavior plays crucial roles in chromosome capture, alignment at the metaphase plate, spindle positioning in the cell, and chromosome segregation during anaphase (Grill et al., 2003; Grill & Hyman, 2005; Maiato et al., 2017; Steblyanko et al., 2020; Stumpff et al., 2012). In cultured mammalian cells, chromosome segregation in anaphase is accomplished as kinetochore fibers shorten, moving chromosomes closer to the spindle poles (anaphase A) and by spindle elongation (anaphase B) which is accompanied by elongation of overlapping antiparallel midzone microtubules (Brust-Mascher & Scholey, 2011; Maiato & Lince-Faria, 2010; McIntosh et al., 2012; Roostalu et al., 2010; Vukušić & Tolić, 2021). Midzone microtubules, also called bridging fibers, are nucleated in the half-spindle in an augmin dependent manner and link sister kinetochore fibers contributing to both chromosome alignment and segregation (Jagrić et al., 2021; Štimac et al., 2022; Uehara et al., 2009, 2016). How midzone microtubules contribute to force generation in anaphase varies in different cell types (Vukušić & Tolić, 2021). For example, in meiotic spindles, midzone microtubules generate pushing forces that drive chromosome segregation (Laband et al., 2017). The contribution of midzone microtubule polymerization and sliding has also been documented in yeast and diatom spindles (Cande & McDonald, 1985; Janson et al., 2007; Khodjakov et al., 2004; Vukušić & Tolić, 2021). In cultured mammalian cells, force generation by the midzone changes over time, with pushing forces seen in early and braking forces in later anaphase (Collins et al., 2014; Pamula et al., 2019; Saunders et al., 2007; Vukušić et al., 2017; Yu et al., 2019). In some cell types, interactions of astral microtubules with cortical force generators contribute to spindle elongation in anaphase (Aist et al., 1991, 1993).

In addition to contributing to chromosome segregation, midzone microtubules play a well-established role in specification of the location of contractile ring formation (Alsop & Zhang, 2003, 2004; Cao & Wang, 1996; Wheatley, 1996). Several protein complexes, including the chromosome passenger complex and centralspindlin, accumulate on midzone and equatorial astral microtubules and generate signals that activate the GTPase RhoA at the equatorial cell cortex (Adriaans et al., 2019; Canman et al., 2008; Hirose et al., 2001; Hutterer et al., 2009; Jantsch-Plunger et al., 2000; Mollinari et al., 2002; Nishimura & Yonemura, 2006). RhoA in turn activates formins and myosin to promote contractile ring formation and constriction (Li & Higgs, 2003; Matsumura, 2005).

A key component of midzone formation and function is the conserved antiparallel microtubule crosslinking protein, PRC1 (Kurasawa et al., 2004; Mollinari et al., 2002; Vernì et al., 2004; Zhu & Jiang, 2005). Mammalian cells lacking PRC1 fail to generate robust midzone arrays and frequently fail cytokinesis (Adriaans et al., 2019; Kurasawa et al., 2004; Mollinari et al., 2005). PRC1 interacts with several midzone components, including the microtubule plus-TIP binding protein CLASP1 which promotes microtubule growth (Aher et al., 2018; Al-Bassam et al., 2010; Liu et al., 2009; Maton et al., 2015). PRC1 also interacts with the centralspindlin subunit MKLP1 - an interaction which contributes to the mechanical properties of the midzone (Hutterer et al., 2009; Lee et al., 2015; Mishima & Lee, 2015). PRC1 regulates midzone length by recruiting the kinesin Kif4a, which suppresses the elongation of midzone microtubule plus-ends (Hu et al., 2011; Nunes Bastos et al., 2013; Subramanian et al., 2013). Live cell imaging of tagged PRC1 further shows that as anaphase progresses, PRC1 intensity increases and dynamic turnover decreases (Asthana et al., 2021; Pamula et al., 2019). The change in the extent of microtubule overlap measured in live cells expressing fluorescent PRC1 (Asthana et al., 2021; Pamula et al., 2019) confirms earlier results obtained by EM (Mastronarde et al., 1993).

*In vitro* experiments with purified proteins provide additional insight into how PRC1 and Kif4a regulate antiparallel microtubule dynamics. On individual microtubules PRC1 and Kif4a accumulate forming dynamic end tags; the length of the tag is proportional to PRC1 concentration and microtubule length (Subramanian et al., 2010, 2013). Mixtures of PRC1, Kif4a and microtubules also form antiparallel bundles *in vitro* (Bieling et al., 2010; Hannabuss et al., 2019; Kapitein et al., 2008; Wijeratne & Subramanian, 2018). Microtubule overlap zones shorten as microtubules slide apart and eventually reach steady state; and overlap length is similar to the length of midzone overlap zones observed in cells (Bieling et al., 2010; Hannabuss et al., 2019). PRC1 has been shown to generate frictional forces to resist microtubule sliding, consistent with the braking function of midzones observed in cells (Aist et al., 1991, 1993; Gaska et al., 2020; Pamula et al., 2019; Saunders et al., 2007). Modeling shows that microtubule sliding results in condensed regions of PRC1 near microtubule plus-ends (Hannabuss et al., 2019).

Despite information about the dynamics and distribution of PRC1 in midzones, the dynamic behavior of the microtubules to which PRC1, and other midzone proteins, are bound has not been systematically measured throughout anaphase. Here we report the results of experiments in live cells showing that midzone microtubules initially comprise a single population of highly dynamic microtubules and subsequently transition to a mixture of dynamic and more stable microtubules. By mid-cytokinesis, these microtubules are converted to a static array for which no detectable turnover/dissipation can be measured. The rate of microtubule sliding from early anaphase to late anaphase, as antiparallel overlap length decreases. Conversion to a static array occurs when highly compacted zones of PRC1 are measured in cells. Prevention of lateral compaction, by blocking cytokinesis, or longitudinal compaction by depletion of Kif4a or PRC1, results in midzone microtubule arrays that fail to form static arrays.

## Results

### Gradual Reduction in Midzone Microtubule Turnover and Sliding in Anaphase

To understand microtubule dynamics throughout anaphase, we first quantified chromosome and pole motion during anaphase in LLCPk1 pig epithelial cells expressing EGFP-α-tubulin. Consistent with prior work in these cells, chromosomes move closer to the spindle poles as kinetochore fibers shorten (anaphase A) and simultaneously the distance between the spindle poles increases as midzone microtubules elongate (anaphase B) (Figure 1A) (Collins et al., 2014; Rusan & Wadsworth, 2005). The rate of chromosome motion is faster within the first minute of anaphase onset (AO) and gradually slows after 5 minutes (avg. rates of 2.4 and 1 µm/min, respectively). Similarly, the rate of pole-to-pole separation is rapid within the first minute and decreases after 5 minutes (avg. rates of 1.8 and 0.5 µm/min, respectively). To quantify midzone microtubule dynamics throughout anaphase, we photoactivated cells stably expressing PA-EGFP-α-tubulin and quantified the dissipation of fluorescence and the position of the marked region (Tulu et al., 2003, 2010). In preliminary experiments, we photoactivated cells twice, first in early anaphase (Figure 1B, top) and then again between 4 - 6 minutes following the first activation (Figure 1B, bottom). In each of the six experiments, the half-time for fluorescence dissipation increased (Figure 1C, top) and the velocity of microtubule sliding decreased from the first to second activation (Figure 1C, bottom) confirming work in other cell types demonstrating that microtubule dynamics and sliding are rapid in early anaphase (Vukušić et al., 2017; Yu et al., 2019) and are slower in late anaphase and telophase (Pamula et al., 2019). For these experiments images were collected at a 10 - 30 second interval and data were best fit with a single exponential (see Methods). This relatively long interval between images will not capture rapid events, so we next performed single activations at various times following AO using a 5 second interval between images.

**Figure 1.**
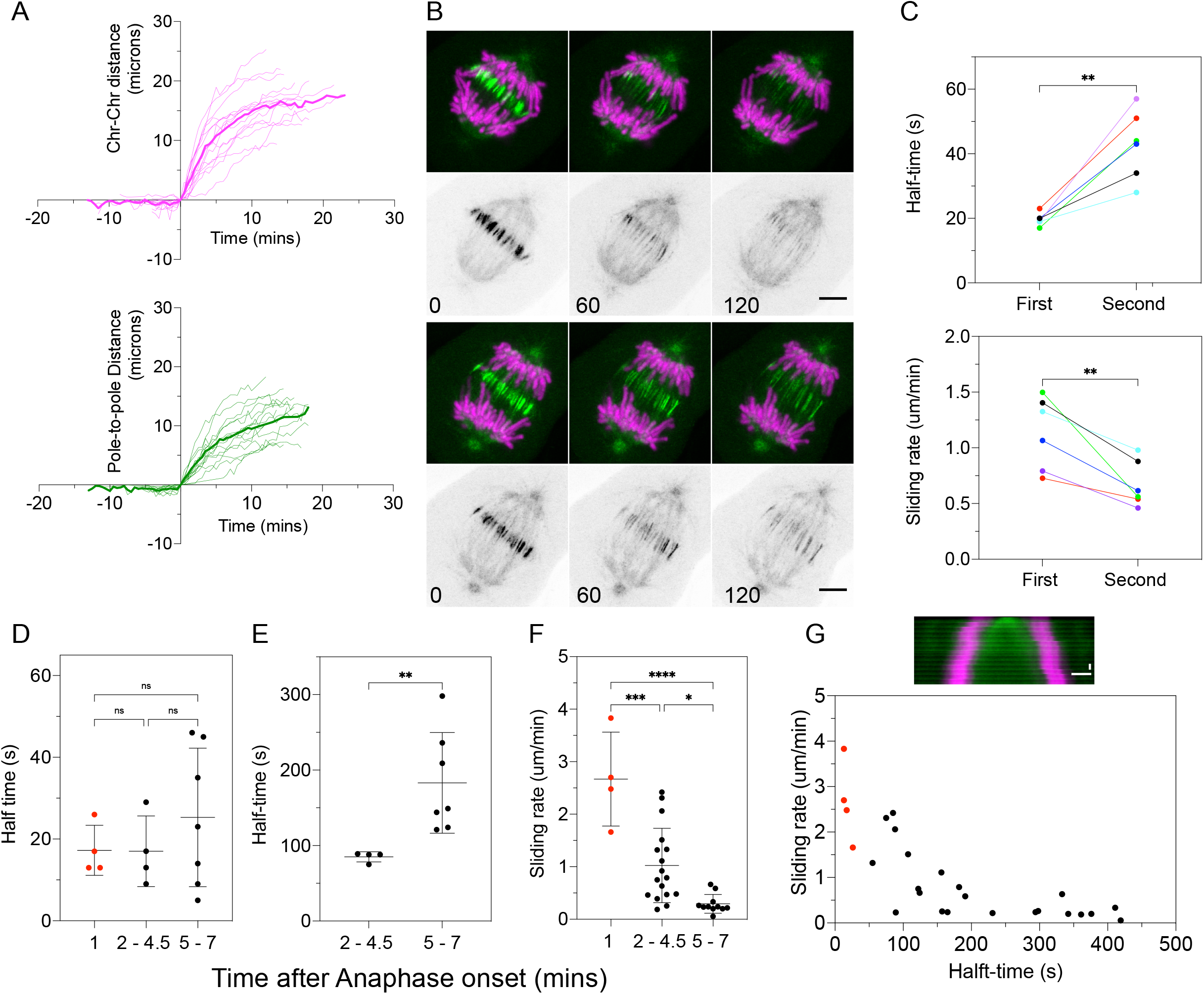
Gradual reduction in midzone microtubule turnover and sliding in anaphase. (A) Chromosome-chromosome (top) and pole-to-pole (bottom) segregation in anaphase (dark color, average; pale color, individual cells; n = 15 cells; Time 0 is anaphase onset (AO)). (B) Double photoactivation of a representative anaphase cell showing first activation (top 2 rows) and second activation (bottom 2 rows). Top rows: Photoactivated microtubules (green) and DNA (magenta). Bottom rows: microtubules in grayscale. Bar = 5 µm. Time in seconds. (C) Plot of dissipation half-time (top) and sliding rate (bottom) for the first and second activations. Each color is an individual cell (n = 6 cells). Plots of fluorescence dissipation (D, E) and sliding rate (F) for individual cells activated once. Dissipation half-times for fast (D) and slow (E) populations in individual cells activated once. D, E, n = 14 cells; F, n = 32 cells; x-axis is time after AO. (G) Representative kymograph showing microtubule sliding following photoactivation; chromosomes magenta and photoactivation zone green. Vertical bar = 60 seconds; horizontal bar = 5 µm. Plot of dissipation half-time for the more stable population vs sliding rate; each dot is a single cell. Red dots are cells with only a single, fast half-time. Statistical analysis (C) Paired T-test; (D and F) ANOVA with the post-hoc Tukey test; (E) Mann-Whitney U test; p-values: ns = P > 0.12; * = P < 0.05; ** = p < 0.0021; *** = P < 0.0002; **** = P < 0.0001***.

When activations were performed within approximately 1 min of AO, fluorescence dissipation was best fit by a single exponential with an average half-time of dissipation of 17± 6s (mean± SD, n = 4 cells) (Figure 1D). When activations were performed during mid-and late anaphase (2 - 4.5 and 5- 7 min post AO, respectively), however, the data were best fit to a double exponential resulting in two half-times (see Methods), indicative of two populations of microtubules with different dynamics. The fast phase had an average half-time that was not different from the half-time measured in the early anaphase cells (17± 9s and 25± 17s) for mid-and late anaphase respectively (Figure 1D). The less dynamic population of microtubules had an average half-time of 85± 9s in mid anaphase which increased to an average of 183± 67s in late anaphase (Figure 1E).

As the midzone elongates, overlapping, antiparallel microtubules slide relative to each other. To measure the dynamics of sliding, we quantified the distance that photoactivated marks on midzone microtubules moved (see Methods). The rate of sliding decreased from an average of 3± 0.9 µm/min in early anaphase to 1± 0.7 and 0.3± 0.2 µm/min in mid and late anaphase cells, respectively (Figure 1F) (Pamula et al., 2019). Comparison of sliding and turnover also showed that microtubules in early anaphase, which show rapid turnover, slide faster than microtubules in late anaphase (Figure 1G). In addition, the average rate of microtubule sliding in early anaphase is similar to the rate of chromosomes but faster than the rate of pole-to-pole separation as observed in human cells (Supplemental Figure 1B) (Pamula et al., 2019; Vukušić et al., 2017).

### Midzone Microtubule Turnover during Contractile Ring Ingression

Midzone microtubules provide signals which promote the assembly of an actomyosin ring at the equatorial cell cortex. Contraction of the ring causes ingression of a cleavage furrow which constrains components of the midzone into a focused structure called the midbody (Hu et al., 2012; Peterman & Prekeris, 2019).

To examine midzone microtubule dynamics during cytokinesis in LLCPk1 cells, we first quantified furrowing in these cells. Cytokinetic ingression begins at an average of 9± 1 minutes after AO, and the furrow is approximately 50% ingressed by an average of 12± 2 minutes after AO (Figure 2A). Note that in these epithelial cells cleavage is asymmetric with the basal side completing furrowing prior to the apical side where cell-cell junctions are present, so the extent of furrowing was measured in the plane of the spindle toward the apical side (Higashi et al., 2016).

**Figure 2.**
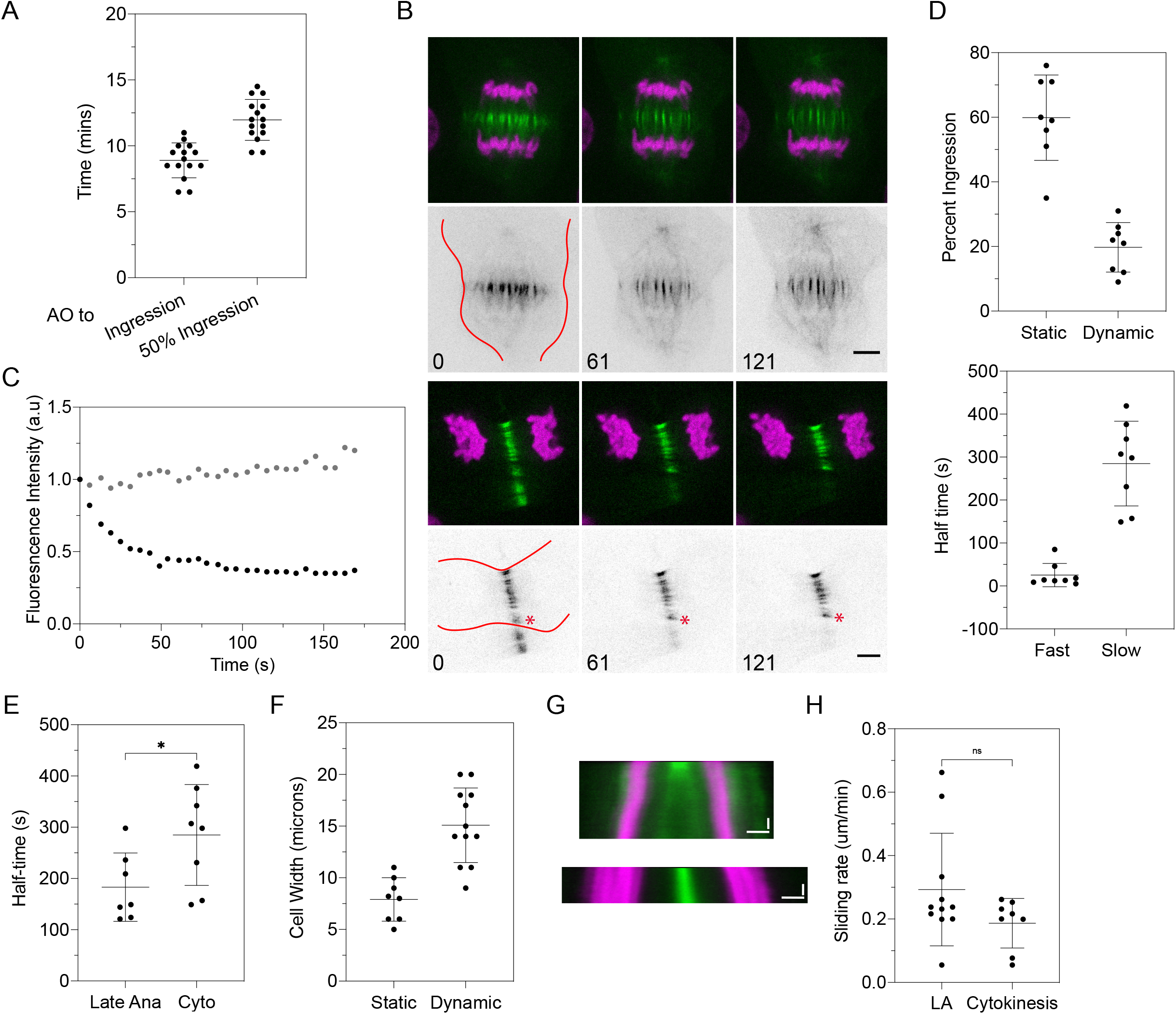
Microtubules transition from stable to static at mid-cytokinesis. (A) Time from AO to the onset of ingression and to 50% ingression (n = 15 cells). (B) Photoactivation of midzone microtubules at onset (top cell) and following 50% ingression (lower cell). Top rows: Photoactivated microtubules (green) and DNA (magenta). Bottom rows: microtubules in grayscale. Bars = 5 µm. Cell edge outlined in red in first frame; red asterisk marks peripheral bundle. Time in seconds. (C) Dissipation curves for representative cells photoactivated at onset (black) and after 50% ingression (gray). AU; arbitrary units of fluorescence; time in seconds. (D) (Top) cells were scored for dynamic or static microtubules and percentage of ingression was measured (n = 16 cells); (Bottom) half-times of fluorescence dissipation for cytokinetic cells with dynamic microtubules (n = 8 cells). (E) Comparison of dissipation half-time for the more stable population in late anaphase and cytokinesis. (F) Cells were scored for dynamic or static microtubules and cell width at the equator was measured (n = 20 cells). (G) Kymographs of chromosomes and photoactivated microtubules in cells photoactivated at the onset (top) and following 50% ingression (bottom); DNA in magenta, photoactivation zone in green. Vertical bar, time = 60 seconds; horizontal bar = 5 µm. (H) Plot of rate of microtubule sliding for cytokinetic cells at onset of ingression (n=8 cells); late anaphase (LA) shown for comparison. Statistical analysis (E) Mann-Whitney U test; p-values: * = P < 0.05.

To investigate microtubule turnover during cytokinesis, midzone microtubules were photoactivated and the percentage of ingression and the cell width at the equator were determined for each cell. As shown in Figure 2, midzone activations performed prior to ∼50% ingression had measurable fluorescence dissipation and were categorized as dynamic (Figure 2B, top and 2C, black trace). In contrast, for cells with greater than ∼50% ingression, a half-time could not be determined from the dissipation curve; microtubules in these cells were classified as static (Figure 2B, bottom and 2C, gray trace). For cells with less than ∼50% ingression, the dissipation curve was best fit to a double exponential (Figure 2C, black trace). The half-time for the fast population was not different from that measured in cells in early anaphase (avg. half-time of 24± 30s, n = 6 cells; see Figure 1 and Supplemental Figure 1C, left). The half-time for the more stable microtubule population, however, was greater than that measured in late anaphase cells, demonstrating the continued stabilization of midzone microtubules over time (avg. half-time of 305± 97s, n = 6 cells; Figure 2D, bottom and 2E). Similar results were obtained when cell width was used to estimate the extent of ingression (Figure 2F). This data shows that the transition to a static array occurred when cell width is less than approximately 8 µm.

Midzone microtubule sliding was also measured during cytokinesis (Figure 2G - H). Cells at less than ∼50% ingression showed sliding at an average rate of 0.2± 0.07 µm/min, which was not statistically different from sliding in late anaphase cells (Figure 2H). Sliding could not be detected in cells with greater than ∼50% ingression (Figure 2G).

### PRC1 and Kif4a regulate midzone microtubule turnover

Midzone formation requires the conserved microtubule binding protein PRC1 (Kurasawa et al., 2004; Mollinari et al., 2005; Verbrugghe & White, 2004; Zhu & Jiang, 2005). In metaphase and early anaphase PRC1 localizes to long stretches of antiparallel microtubules and is dynamic as measured by FRAP (Asthana et al., 2021; Polak et al., 2017). As anaphase progresses PRC1 and its binding partner Kif4a localize to short zones at the equator of the midzone (Asthana et al., 2021; Kurasawa et al., 2004; Mollinari et al., 2005; Pamula et al., 2019). This dynamic reorganization of PRC1/Kif4a ultimately acts to prevent further midzone elongation (Kurasawa et al., 2004; Pamula et al., 2019). To determine if and how these midzone proteins impact midzone microtubule dynamics, we depleted each protein using siRNA (Methods; Supplemental Figure 2).

Kif4a and the centralspindlin subunit, MKLP1, were not detected in cells depleted of PRC1 (Supplemental Figure 2B, top panels), confirming the requirement for PRC1 to generate a robust midzone (Adriaans et al., 2019; Kurasawa et al., 2004; Pamula et al., 2019).

We next examined microtubule dynamics in anaphase and telophase cells following treatment with siRNA targeting PRC1. In anaphase cells activated 3 - 6 min post AO, we observed a single population of highly dynamic microtubules. The average half-time for these cells, 27± 10s (n=4 cells), was not different from that measured in control early anaphase cells (Supplemental Figure 2E). These data demonstrate that midzone microtubules in anaphase cells depleted of PRC1 remain as dynamic as microtubules in control cells in early anaphase.

Next, we tested the contribution of PRC1 to midzone microtubule dynamics in telophase cells at >50% ingression, a time when midzone microtubules in control cells have converted to a static array. In telophase cells, we can also confirm that each cell is depleted by measuring the distance between the two sets of segregating chromosomes (Supplemental Figure 2B, left; see Methods). In contrast to control cells, midzone microtubules in telophase cells depleted of PRC1 remained dynamic and the dissipation curve was best fit by a double exponential (Figure 3A). The faster population (average dissipation half-time 16± 12s, n = 11 cells; Figure 3C) was indistinguishable from the fast population of microtubules in control anaphase cells (Figure 1 and Supplemental Figure 1C, left). The slower population had an average half-time of 121± 72s n = 11 cells, which is like that observed in control cells during mid-anaphase (Supplemental Figure 1C, right). Sliding was not measured in these cells because the reduced number of microtubules in the midzone precluded accurate measurements.

**Figure 3.**
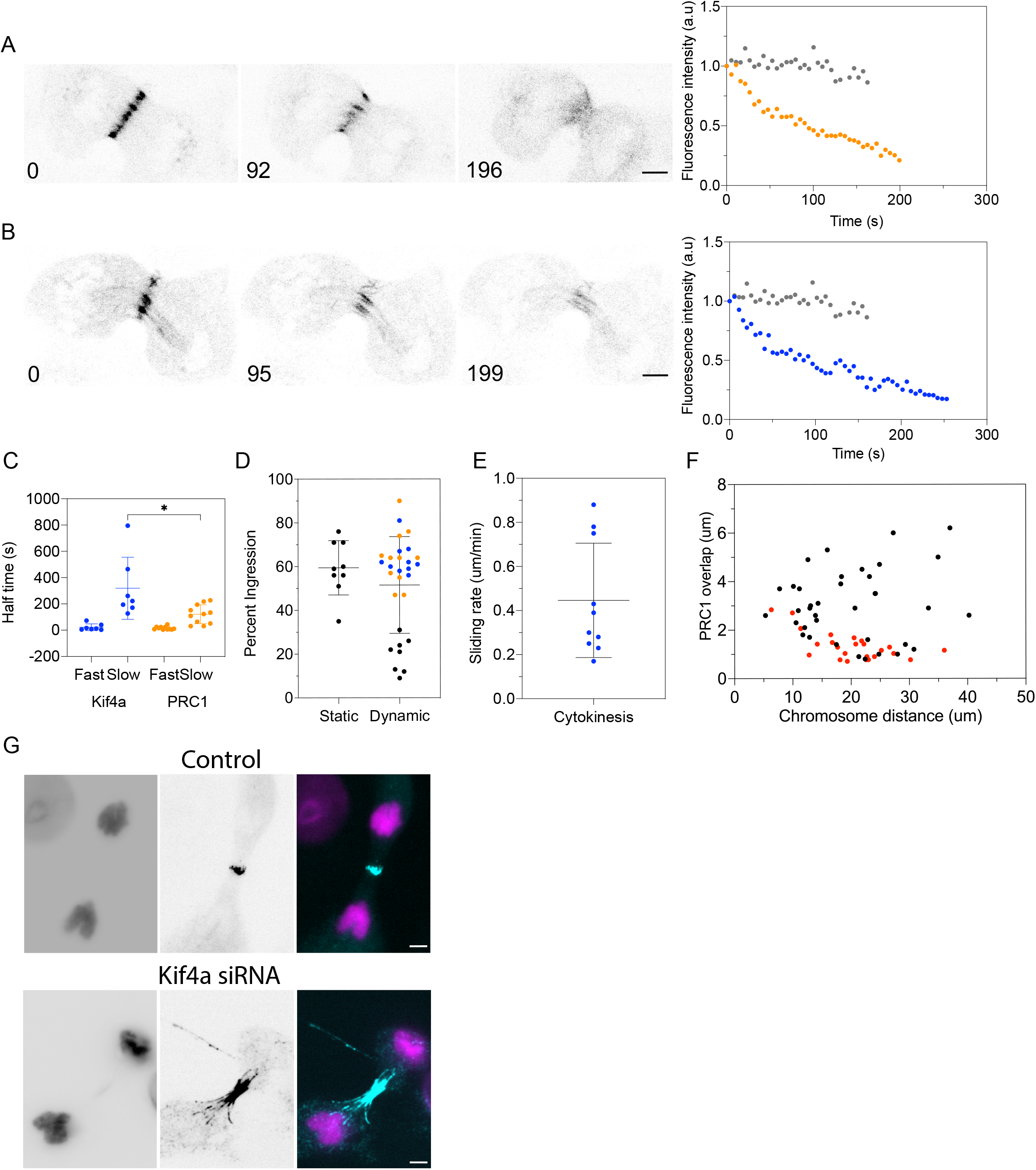
PRC1 and Kif4a impact microtubule stabilization in anaphase and telophase. Photoactivation following 50% ingression in cells depleted of PRC1 (A) or Kif4a (B). Bar = 5 µm. Time in seconds. (Right panels) Dissipation curves for representative cells after 50% ingression, PRC1 (orange), and Kif4a depleted cells (blue). Dissipation for control telophase cell (gray) for comparison. (C) Plot of dissipation half-times for Kif4a and PRC1 depleted cells in telophase (Kif4a n = 7 cells; PRC1 n = 11 cells). (D) Cells were scored for dynamic or static microtubules and percentage of ingression for control (black), Kif4a (blue) and PRC1 (orange) depleted cells (control n = 17 cells; Kif4a n = 10 cells; PRC1 n = 11 cells; control cells from Figure 2E. (E) Plot of rate of midzone sliding for Kif4a depleted cells at > 50% ingression (n= 10 cells). (F) Plot of length of PRC1 overlap zone for control (red) and Kif4a depleted cells (black); Chr-Chr distance on x-axis. (G) Immunofluorescence staining for PRC1 in control (top) and Kif4a depleted cells (bottom); left DNA; middle PRC1; right merge with DNA in magenta and PRC1 in blue. Bar = 5 µm. Statistical analysis (C) Mann-Whitney U test; p-values: * = P < 0.05.

### Kif4a depleted cells

One mechanism by which loss of PRC1 could impact midzone dynamics is through its binding partner, Kif4a, which suppresses microtubule plus-end elongation. Kif4a recruitment to the midzone by PRC1 regulates the length of antiparallel midzone microtubules both *in vitro* and *in vivo* (Bieling et al., 2010; Hannabuss et al., 2019; Hu et al., 2011). Imaging of EGFP-α-tubulin expressing LLCPk1 cells depleted of Kif4a showed that midzone microtubules were frequently curved or buckled (Supplemental Figure 2A, right); these microtubules were longer and the distance between nuclei in telophase was greater than in control cells (Supplemental Figure 2B, right). Microtubule bundles in Kif4a depleted cells appeared less robust than in control cells consistent with previous electron microscopy observations (Hu et al., 2011) (Supplemental Figure 2A, right). Staining Kif4a depleted cells for PRC1 demonstrates that antiparallel microtubule overlap length is longer than observed in control cells (Figure 3F - G). The majority of overlaps are likely composed of antiparallel microtubules; however, it has been shown that PRC1 can also decorate individual microtubules (Subramanian et al., 2013).

First, we measured microtubule dynamics in Kif4a depleted cells in anaphase. Given the variability of microtubule dynamics as anaphase progresses, we used sequential photoactivations in individual cells (see Figure 1B - C), an approach that reveals if microtubule dynamics changes as cells progress through anaphase. Similar to the control cells, Kif4a depleted cells were photoactivated twice, first in early anaphase and then again between 3 - 6 minutes following the first activation. In each of the control cells, microtubule turnover in anaphase slowed from the first to second activation; in Kif4a depleted cells, however, turnover was similar for the first and second activations (Supplemental Figure 2D). The rate of microtubule sliding was also measured in these cells. For control cells, the rate of sliding was faster in early anaphase and then decreased. However, the rate of sliding for Kif4a depleted cells was not significantly different between the first and second activations (Supplemental Figure 2D) (Vukušić et al., 2021). These data show that the gradual reduction in turnover observed in control cells was not detected in Kif4a depleted cells. In addition, the rate of microtubule sliding was generally faster consistent with the retention of longer overlaps in the depleted cells (Figure 3F - G) (Hu et al., 2011; Pamula et al., 2019).

Next, we performed photoactivations in Kif4a depleted cells during telophase. Dissipation curves were best fit with a double exponential with an average half-time for fast population of 23± 27s which was not different from the dynamic population in control cells during anaphase and early cytokinesis (Supplemental Figure 1C). The dissipation half-time for the more stable microtubules in Kif4a depleted cells (319± 236s, n= 7 cells) was significantly greater than in PRC1 depleted cells (121± 72s), likely due to the retention of PRC1 in Kif4a depleted cells (Figure 3B - G). Thus, as observed in PRC1 depleted cells, static microtubule arrays failed to form in telophase cells lacking Kif4a (Figure 3B). Microtubule sliding in telophase cells depleted of Kif4a was like that observed in control cells during mid-to late anaphase (Figure 1F and 3E).

### Contractile Ring Ingression impacts Microtubule Behavior in Telophase

Our results demonstrate that midzone microtubules are gradually stabilized and ultimately rendered static as cells exit mitosis. In addition, our data show that formation of a narrow, tightly focused zone of microtubule overlap, marked by PRC1, contributes to microtubule stabilization in anaphase and is required for the formation of a static midzone array in telophase. Previous work showed that stabilization of midzone microtubules, measured by resistance to cold or nocodazole induced disassembly, required contractile ring ingression (Cimini et al., 1998; Landino & Ohi, 2016). Our results demonstrate that when PRC1 is depleted, static arrays are not formed despite ingression. To explore if and how constriction of the contractile ring impacts midzone dynamics, we inhibited contractile ring formation using latrunculin B (LatB) which binds and sequesters actin monomer (Fujiwara et al., 2018; Spector et al., 1983) or inhibited Rho-GTPase with C3 transferase to prevent activation of myosin and formins (Aktories et al., 2005; Kanada et al., 2009). Cells treated with LatB or C3 failed to ingress and an F-actin-containing contractile ring did not assemble at the equatorial cortex (Figure 4A) (Kanada et al., 2009). Microtubule distribution in LatB or C3 treated cells revealed bundled microtubules in the center of the midzone like control cells (Figure 4A; see also Figure 3H). In addition, a second population of microtubules, which we refer to as peripheral microtubules, was detected in the treated cells (Kanada et al., 2009). These microtubules were located between the non-ingressed membrane and the more central region of the midzone; additionally, prominent astral arrays were present in most treated cells (not shown) (Kanada et al., 2009; Verma & Maresca, 2019).

**Figure 4.**
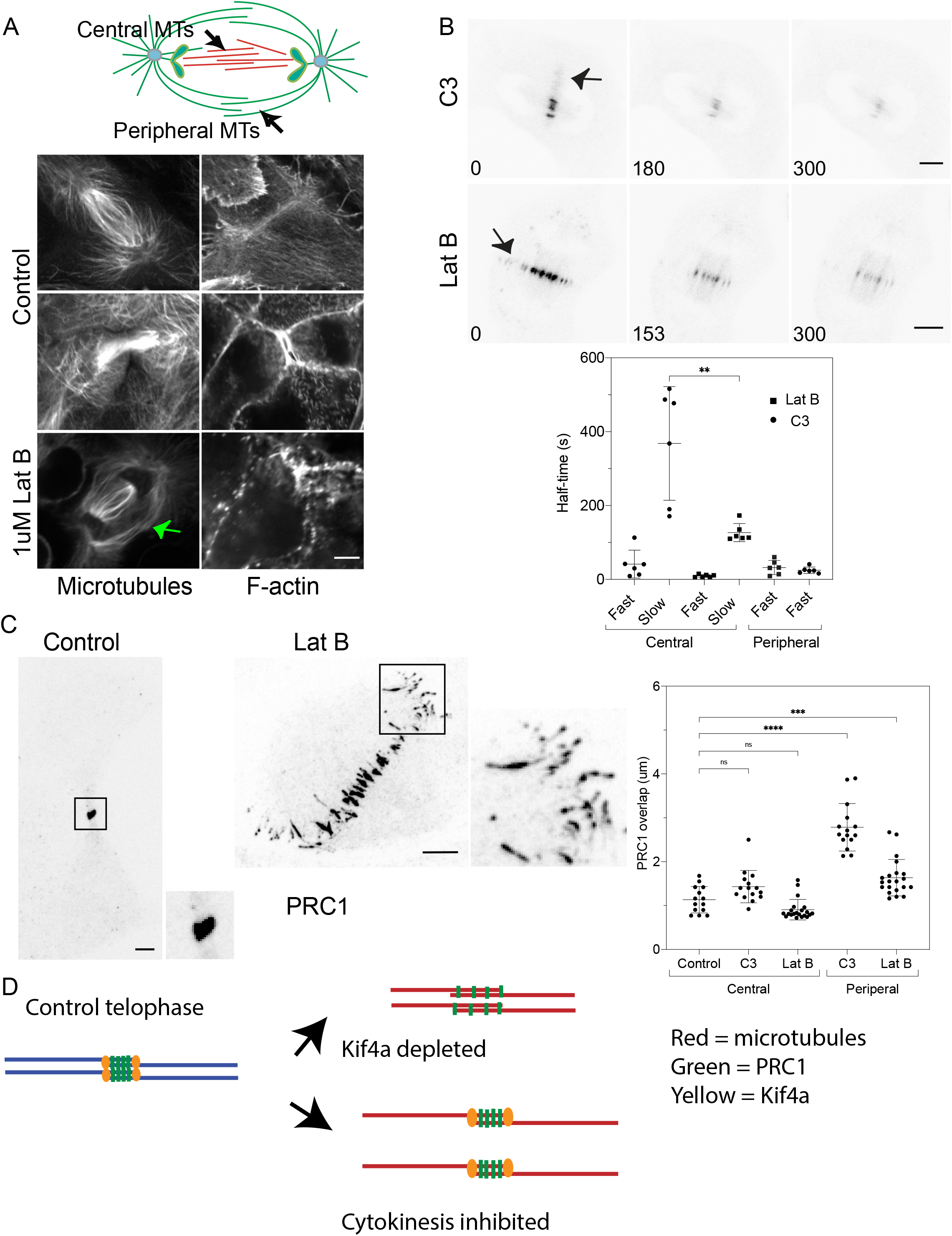
Transition to static microtubules requires lateral compaction of midzone microtubules at cytokinesis. (A) Diagram (top) showing peripheral and central midzone microtubules; (bottom) representative cells showing distribution of microtubules and F-actin in control cells in early and late cytokinesis (top two rows) and following treatment with Latrunculin B (bottom). Left panels microtubules; right panels F-actin. Peripheral microtubules marked with green arrow. (B) Photoactivation of midzone microtubule in a representative telophase cell treated with C3 (top) or Latrunculin B (bottom). Bar = 5 µm. Time in seconds. Peripheral microtubules marked with black arrows. (Lower) Plot of fluorescence dissipation half-times for central and peripheral microtubules in C3 and Latrunculin B treated cells. (C) Immunofluorescence of PRC1 in the midzone of control cells (left) and Latrunculin B treated cells (right) and graph of length of PRC1 for peripheral and central microtubules. Higher magnification views of the region enclosed by the black box are shown on the right of control and bottom of the treated cell. (D) Diagram showing compaction of PRC1 in telophase control, kif4a depleted cell (top), and failed regression (bottom). Statistical analysis (B) Mann-Whitney U test; (C) ANOVA with the post-hoc Tukey test; p-values: ** = p < 0.0021; *** = P < 0.0002; **** = P < 0.0001.

For these experiments, cells were photoactivated in telophase, as marked by nuclear reformation, to ensure that cytokinesis had been inhibited. The results revealed distinct kinetics for central and peripheral microtubules (Figure 4B). The dissipation of fluorescence for peripheral microtubules in both C3 and LatB treated cells was best fit with a single exponential, with half-times for dissipation of 32± 19s and 25± 9s for C3 and LatB respectively, which are not different from the fast population of microtubules in control anaphase cells (Figure 4C, Supplemental Figure 1C). In control cells, peripheral microtubules are occasionally observed (Rusan & Wadsworth, 2005) however because these microtubules were of low abundance and present only transiently photoactivation experiments were unsuccessful. Of note, in control cells, peripheral bundles are stabilized during ingression, showing behavior like the more centrally located microtubules (see bundle marked with asterisk in Figure 2B, bottom panel). Together, these results demonstrate that cells blocked in cytokinesis contain a population of peripheral microtubules which have dynamic behavior like microtubules in early anaphase cells (Supplemental Figure 1C).

The dynamic turnover of the central microtubules in cells blocked in cytokinesis was also determined. Fluorescence dissipation curves for central microtubules in LatB and C3 treated cells were best fit with a double exponential with dynamic and more stable populations (Figure 4B, lower). The dynamic population had dissipation half-times like those measured for control cells (see Supplemental Figure 1). The values for the slow population of microtubules in the central region of LatB and C3 treated cells were like early and late anaphase control cells, respectively. The more stable population in C3 treated cells (368± 154s, n=6 cells) was highly variable and statistically different from LatB treated cells (127± 25s, n=6 cells) (Figure 4B). This difference is likely related to the duration of treatment with each inhibitor. Photoactivations in LatB treated cells were performed within 30 minutes of drug addition, because longer incubations resulted in loss of adhesion and cell shape, precluding successful experiments. In contrast, inhibition of RhoA with C3 required longer incubations (see Methods) and cells selected for activation could have been blocked in telophase for up to several hours.

To gain insight into the organization of central and peripheral microtubules, the length of PRC1 zones was measured in C3 and LatB treated cells. For both treatments, PRC1 zones on peripheral microtubules were longer than for central microtubules. The length of PRC1 zones on central microtubules were not different from control telophase cells (Figure 4C, right). Longer PRC1 zones on peripheral microtubules could result from PRC1 decorating individual, non-overlapping microtubules or from longer antiparallel overlap zones. For both central and peripheral microtubules, PRC1 zones did not compact laterally but remained spaced apart (Figure 4C). Our experiments show that constriction of the contractile ring gathers and stabilizes peripheral microtubules in normal cytokinesis (see Figure 2 and 3H) and that in the absence of constriction, microtubules are retained in the peripheral regions, and fail to stabilize.

## Discussion

Our quantitative analysis of midzone microtubule dynamic behavior demonstrates that midzone microtubules transition from a single, highly dynamic population to a mixture of dynamic and more stable microtubules as anaphase progresses. Furthermore, the more stable population becomes even less dynamic over time. These data are consistent with prior observations, using photoactivation or photobleaching that measured the behavior of midzone microtubules in early (Vukušić et al., 2021; Yu et al., 2019) and late anaphase and telophase (Saxton & McIntosh, 1987). The observation that the more stable set of midzone microtubules became more stable is consistent with time dependent modulation of microtubule dynamics as cells exit mitosis (Mollinari et al., 2002; Neef et al., 2007). Previous work showed that midzone microtubules are generated by augmin-mediated nucleation in the half-spindle and midzone regions (Goshima et al., 2008; Štimac et al., 2022; Trupinić et al., 2022; Uehara et al., 2016).

Our results showing that new, dynamic microtubules are present throughout anaphase suggests that the augmin-dependent mechanism continually generates microtubules, as the half-spindle shortens, and cells exit mitosis (Johmura et al., 2011; Timón Pérez et al., 2022; Tsai et al., 2011; Zhang et al., 2022). The final transition, from a mixture of stable and dynamic microtubules to a static array, occurred when cells reached approximately 50% ingression or a width of ∼8um. Our results expand on prior work that used sensitivity to nocodazole or cold treatment to assess midzone microtubule stabilization (Landino & Ohi, 2016). The prior results showed that microtubules in post-ingression but not in pre-ingression cells were resistant to disassembly, consistent with time-dependent stabilization (Hu et al., 2011; Landino & Ohi, 2016). These experiments also showed that blocking ingression with either blebbistatin or cytochalasin prevented stabilization (Landino & Ohi, 2016). In contrast, we observed that microtubules became stable when furrowing was blocked, but the final transition to a static array did not occur. One possible reason for this discrepancy is due to the greater sensitivity of photoactivation which can provide information regarding the extent of stabilization that is not revealed with high doses of nocodazole. Together our experiments and prior work demonstrate that across different cultured cells and in *C*.*elegans* midzone microtubules are gradually stabilized and then abruptly rendered static at approximately 50% ingression (Saxton & McIntosh, 1987; Yu et al., 2019).

The results of these experiments support a model in which both midzone proteins and cytokinesis drive the formation of a static array in telophase cells. Cells depleted for PRC1 or Kif4a fail to form static arrays in telophase despite ingression and the concomitant formation of a bundle of microtubules. Further, these cells lack or have longitudinally expanded PRC1 zones. Conversely, cells that fail to ingress form stable, but not static, midzone microtubules that do not compact laterally. Although static arrays did not form, both Kif4a and PRC1 depleted cells did contain stable microtubules in late anaphase and telophase. These microtubule populations could arise due to incomplete depletion and/or additional midzone components that bundle microtubules. Candidates for microtubule stabilization in the absence of PRC1 include CPC, CLASP1/2, and centralspindlin (Bratman & Chang, 2007; Davies et al., 2015; Maiato et al., 2003). Finally, in cells lacking PRC1, microtubule compaction may promote stability via contractile ring constriction (Hirsch et al., 2022; Verma & Maresca, 2019).

In anaphase, both midzone and astral microtubules contribute signals for cytokinesis, although the former source has generally been considered more important in cultured mammalian cells (Cao & Wang, 1996; Wheatley, 1996). We observed a highly dynamic peripheral array of microtubules in cells blocked in cytokinesis. Control cells also contained peripheral microtubules which were stabilized during furrow ingression (Verma & Maresca, 2019). These microtubules likely arise from nucleation at the centrosome, as astral arrays extending away from the equatorial region were also enhanced in these cells. These microtubules may arise due to the increase in microtubule elongation as cells exit mitosis (Rusan & Wadsworth, 2005). Recent work demonstrates that in *C. elegans* cells lacking a midzone, peripheral microtubules can form a midbody via contractile ring constriction (Hirsch et al., 2022). Our experiments support the observation that multiple mechanisms that can generate microtubules for midzone and midbody formation in telophase (Hirsch et al., 2022).

Taken together our results show that formation of a static array in telophase requires both longitudinal and lateral compaction of PRC1 on microtubules. Our results confirm that PRC1 plays a critical role in midzone formation in mammalian cells and reveal how cytokinesis impacts midzone microtubules, converting a stable array into a static structure.

## Materials

Unless otherwise stated, all reagents were obtained from Sigma-Aldrich.

## Methods

### Cell culture

Experiments were performed on LLC-PK1 pig epithelial cells obtained from the ATCC and tested for mycoplasma. Cells were cultured in a 1:1 mixture of F-10 Hams and Opti-MEM (ThermoFisher) containing 7.5% Fetal bovine serum (CPS Serum FBS-500-HI) and antibiotic-antimycotic. Cells were grown at 37°C and 5% CO2, in a humid atmosphere. In addition to parental cells, experiments were performed on LLC-Pk1 cells expressing EGFP-α tubulin or photoactivatable EGFP-α tubulin generated as previously described (Rusan et al., 2001; Tulu et al., 2003, 2010). For live cell imaging experiments, the growth medium was removed and replaced with non-CO_2_ MEM medium lacking bicarbonate and containing HEPES. A temperature of ∼37°Cwas maintained during imaging using either a Nicholson Precision Instruments Air stream stage incubator (Nicholson Precision Instruments ASI 400) or an Oko stage incubator (20/20 Technologies Inc.). Temperature was monitored using a thermistor probe adjacent to the culture dish (Air stream incubator) or within the heater (Oko).

### Immunofluorescence

LLC-PK1 cells were plated on #1.5 22×22mm coverslips 24-48 hours prior to fixation. Cells were rinsed 2 or 3 times in room temperature calcium and magnesium free PBS, and then fixed in -20°Cmethanol for 4-10 min. Cells were rehydrated in PBS containing 0.1% Tween 20 and 0.02% sodium azide (PBS-Tw-Az). For some antibodies, cells were fixed in freshly prepared 0.25% glutaraldehyde, 2% paraformaldehyde, 0.5% Triton X100, prepared in PBS. Cells were incubated in the manufacturer’s recommended dilution of primary antibody prepared with 2% BSA in PBS-Tw-Az. The following antibodies were used in these experiments: PRC1 (Santa Cruz, 1:40), Kif4a (Santa Cruz,1:500); MKLP1 (Santa Cruz, 1:25), and tubulin (mouse anti-tubulin Dm1α (Sigma) 1:500 or sheep anti-tubulin (Cytoskeleton, 1:250).

Incubations with primary antibodies were performed in a humid chamber for 60 minutes at 37°Cor overnight at room temperature. Coverslips were rinsed by dunking the coverslip at least 30 times in a beaker containing PBS-Tw-Az. Secondary antibody incubations were performed in a humid chamber at room temperature for 45 minutes, at the manufacturers recommended dilution, followed by rinsing as described above.

Coverslips were mounted in a mounting medium (Southern Biotech) containing DAPI and sealed with nail polish. The following secondary antibodies were used: Alexa Fluor 568 goat anti-mouse (Molecular Probes); Cy3 goat anti-rabbit (Jackson ImmunoResearch); Alexa Fluor 488 donkey anti-sheep (Jackson ImmunoResearch). F-Actin was visualized using rhodamine labeled phalloidin according to the manufacturer’s recommended dilution (Molecular Probes). Cells stained with phalloidin were fixed in paraformaldehyde-glutaraldehyde as described above.

### siRNA

For protein depletions, LLC-PK1 cells (parental, or cells expressing EGFP-α tubulin or PA-EGFP-α tubulin) were plated at a density of 250,000 cells per well of a 12 well plate. The following day, cells were incubated with siRNA using Lipofectamine RNAiMax Transfection reagent (Invitrogen) according to the manufacturers’ recommendations. All siRNAs were obtained from Sigma-Aldrich; siRNA was resuspended in RNAi buffer (Dharmacon) to a concentration of 100 uM. The final working concentration was 500 nM for PRC1 siRNA and 250 nM for Kif4a siRNA. Cells were incubated with siRNA for 5 hours. After the 5-hour incubation, cells were trypsinized and replated on either glass bottom dishes (MatTek Corp.) or on glass coverslips. The sequences used for siRNA are: Kif4a 5’GCAGAUUGAAAGCCUAGAG3’; PRC1 5’UUGGGAUUCCAGAGGACCA3’. Transfection efficiency was monitored using SiGLO (Dharmacon); control transfections contained all reagents but no siRNA.

### Inhibitors

C3 transferase was obtained from Cytoskeleton Inc. (CT03) and rehydrated in distilled water to a concentration of 1mg/ml and stored at -80°C. Immediately prior to use, C3 was prepared in a culture medium lacking serum to a final concentration of 4 ug/ml; cells were rinsed in PBS and then incubated with C3 for ∼5 hours at 37°Cin a humid atmosphere.

### Imaging

LLC-PK1 cells expressing EGFP-α tubulin were imaged using a Nikon Ti-E microscope with a CSU-X1 Yokogawa Spinning disc confocal scan head and a 100X/ NA1.4 oil immersion objective. Image acquisition was controlled by Metamorph software (Molecular Devices, LLC). Live cells were imaged with a 30 - 60 second or 2 - 4 minutes interval depending on the experiment. For imaging of fixed cells, complete Z-stacks were collected using a 0.30 μm spacing; exposure time varied depending on the antibody.

Photoactivations were performed on both a Spinning Disc and Resonant Scanning Confocal using Nikon Elements software. Photoactivation was performed after anaphase onset, perpendicular to the spindle long axis, in between separating chromatids, visualized with mCherry H2B or by differential interference contrast optics (DIC). The stimulation type used was a line with a width of 1 pixel.

The scanning confocal system consisted of a Nikon Ti Microscope with A1HD Resonant Scanning Confocal with Galvano scanner and DU4 detector. An APO 60X/1.4NA oil immersion objective and a stage top incubator system (Okolab) was used.

Photoactivation of PA-EGFP-α tubulin, a 405 nm laser was used. For all experiments the duration of pulse was set to 100 ms with no delay with laser power at 10.2%. PA-EGFP fluorescence was imaged using a 488 nm laser at 1.7% laser power, and H2B mCherry chromosomes were visualized using a 561 nm laser at 1.3% laser power. For majority of experiments, images were acquired at 5 second intervals. At each interval, 3 Z-slices at 1 μm spacing were acquired.

The spinning disc system consisted of a Nikon Ti Microscope with a Yokogawa Spinning Disc Confocal and Andor DU-897 camera. An APO VC 100X/1.4 oil immersion objective and stage top incubator system (Okolab) was used. For photoactivation of PA-EGFP-α tubulin, a 405 nm laser at 50% laser power for a duration pulse of 100 ms with no delay was used for all experiments. PA-EGFP fluorescence was imaged using a 488 nm laser with exposure of 300 ms and chromosomes were visualized using a 561 nm laser with exposure of 500 ms. Images were acquired at 10 - 30 second intervals.

Double photoactivation experiments were performed similarly, using a Nikon A1R Resonant Scanning Confocal microscope with an APO 60X/1.4NA oil immersion objective. The stimulation type used was a rectangle with length of 16.6 μm and width of 0.6 μm. For this instrument the following laser settings were used: 405 nm 20.8%, 488 nm laser 2%, and 561 nm laser at 0.5%. Images were acquired every 30 second intervals. At each interval, 3 Z-slices at 1 μm spacing were acquired. Since performing these experiments, the 405 nm laser has been changed on the scanning confocal system.

### Image analysis

To determine the dissipation of fluorescence following photoactivation, images were quantified in FIJI (ImageJ open-source software). An area was drawn around the activated region to measure the fluorescence intensity for each frame. The same region was used for background intensity taken within the spindle. The mean fluorescence intensity values were multiplied by the area to obtain integrative intensity. To determine how much photoactivatable-α tubulin fluorescence was lost, we divided the integrative values of the spindle by the integrative values of the background and subtracted 1 from the ratio. The obtained integrated intensity values of fluorescence dissipation were normalized to the first frame. The normalized intensity values were plotted in Kaleidagraph (Kaleidagraph, Version 4.5 for MacOS. Synergy, Reading, PA, USA.). The plot was fitted to either a single [m1+m2^exp(x/-m3)] or a double [m1*exp(x/-m2) + m3*exp(x/-m4)] exponential. To determine the half-time for each individual cell, the tau value obtained from the fit was multiplied by the ln (2).

The time of anaphase onset for each individual cell was defined as the time point immediately prior to the separation of sister chromatids. For some cells in which the time of AO was not observed, we estimated AO as follows: The average distance between the chromosomes was obtained at 30 second intervals for 10 mitotic cells and divided in two. The data was normalized for anaphase onset at time zero and the data was fit to an exponential rise. The equation y = m2^(1-exp(−m3*x) was simplified to c2 = (−1/m3 * ln (1-c3/m2) and was used to calculate time from distance. We then compared cells with known times of anaphase onset with calculated time and fitted the plot to a simple linear regression curve fit (Supplemental Figure 1A).

### Data Analysis and statistical analysis

Measurements were performed in FIJI and statistical analysis was performed in Prism (GraphPad Prism version 9.4.1 for MacOS, (GraphPad Software, San Diego, California USA, www.graphpad.com).

The length of midzone microtubules in Kif4a-depleted cells was measured on a single z-plane of EGFP-α tubulin anaphase spindles, using the segmented line tool in FIJI. To measure the length of the PRC1 overlap zone, images were analyzed using either Metamorph software or FIJI. Individual slices of complete Z-stacks of stained cells were measured, and the average length of the overlap was determined for each cell. PRC1 overlap was plotted as a function of the length between the segregating chromosomes, measured using the center of the DNA masses. For inhibited cells, only overlap zones in cells with reformed nuclear envelopes were measured.

The time from anaphase onset to onset of furrowing and to 50% ingression was determined from movies of cells expressing EGFP-α tubulin. To quantify cell width and percent of ingression, the width of the cell prior to onset of ingression was measured, or in some cases estimated by fitting a curved line along the sides of the cell. Next the width was measured at the time of activation and the fraction of the initial width to the width at activation was determined. Measurements were performed using FIJI.

Graphs of the data were compiled using Prism (GraphPad Prism version 9.4.1 for MacOS, GraphPad Software, San Diego, California USA, www.graphpad.com). Images were assembled in Adobe Illustrator (Adobe Inc., version 25.4.6. Adobe Illustrator).

## Abbreviations

PA-EGF-: α-tubulin,photoactivable
GFP-tagged: α-tubulin
EM: electron microscopy
PRC1: protein regulator of cytokinesis 1
CLASP1: cytoplasmic linker associated protein 1
MKLP1: mitotic kinesin-like protein 1
Kif4a: kinesin family member 4A

## Acknowledgements

The authors thank Dr. T. Maresca for insightful comments on this work. We thank Dr. J. Chambers for expert assistance with microscopy. Dr. K. Velle provided editorial comments on the manuscript, and advice on Prism software. We thank Ms. Louiza Tizi-Ougdal and Eylen Diaz Soto for excellent assistance in the lab; Ms. Amanda Shorey for collecting preliminary data on protein depletions and experiments with Latrunculin. We especially thank Dr. Heather M. Jordan, a former master’s student, whose work helped pave the way for this study. Funding was obtained from NSF MCB 2134215 and CFR was supported by Soft Materials for Life Sciences National Research Traineeship Program 1545399 and the Spaulding-Smith STEM Fellowship.

## Author contributions

P.W. and J.R. conceived of the project. C.F.d. R., P.W. and J.S. performed the experiments. Y.Z. performed initial photoactivation experiments. C.F.d.R. and J. S. performed data analysis. C.F.d.R. and P.W. assembled the figures. P.W., C.F.d.R. wrote the manuscript with assistance from J.S. and J.L.R.

## Supplemental Figure Legends

**Supplemental Figure 1.**
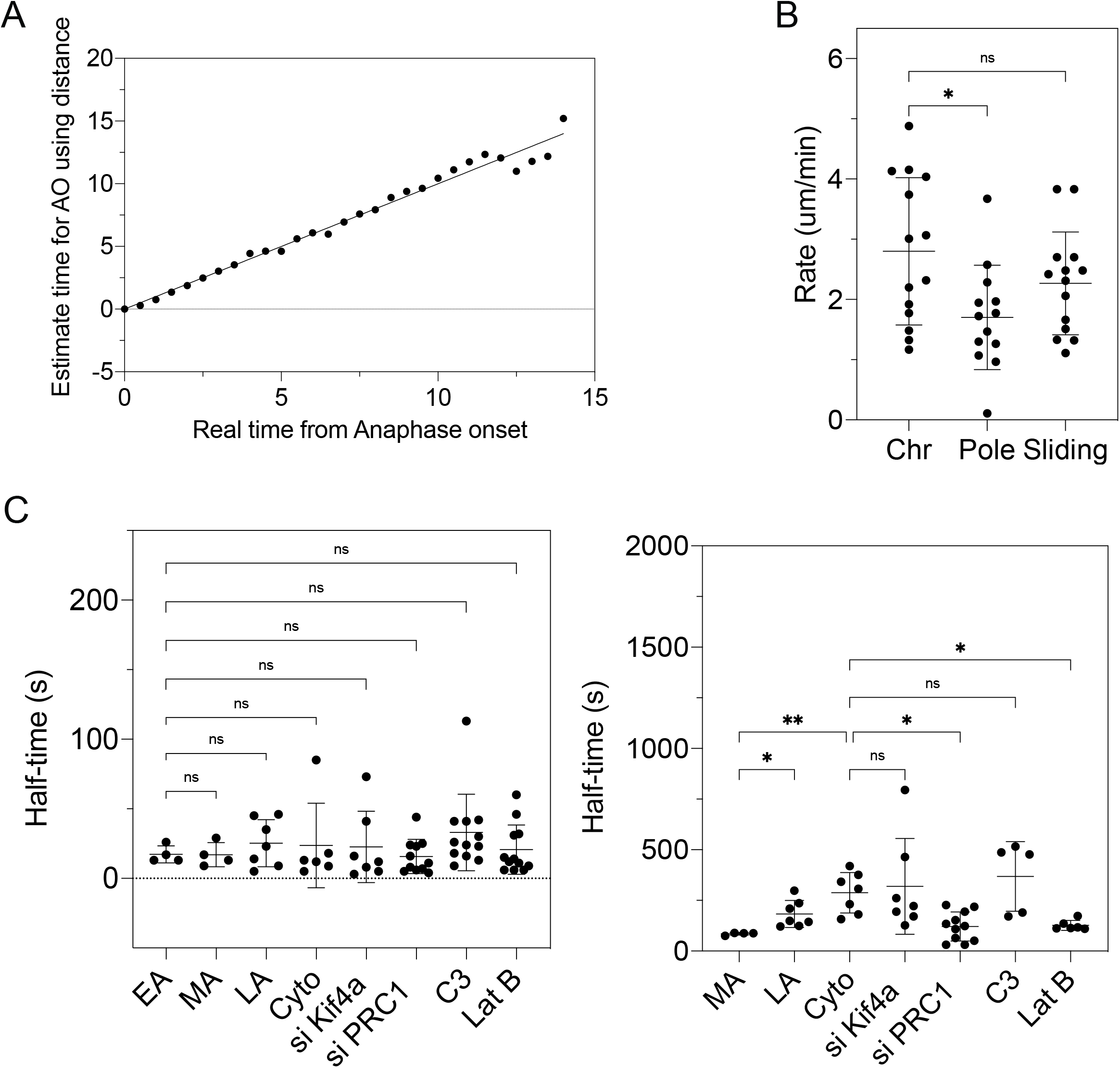
(A) Validation of anaphase onset determination from chromosome-chromosome distance. Plot of the real time from AO vs estimated time for AO from c2 = (−1/m3 * ln (1-c3/m2); Simple linear regression curve fit, R^2^ = 0.9809. (B) Comparison of the rate of chromosome-chromosome, pole-pole and midzone microtubule sliding rates within 2 minutes post AO. (C) Dissipation half-times for the fast population (left) and for the more stable microtubule population (right) for control and treated cells. EA - early anaphase <1 min post AO; MA - mid-anaphase, 2-4.5 min post AO; LA - late anaphase 5-7 min post AO. Statistical analysis ANOVA with the post-hoc Tukey test; p-values: * = P < 0.05; ** = p < 0.0021.

**Supplemental Figure 2.**
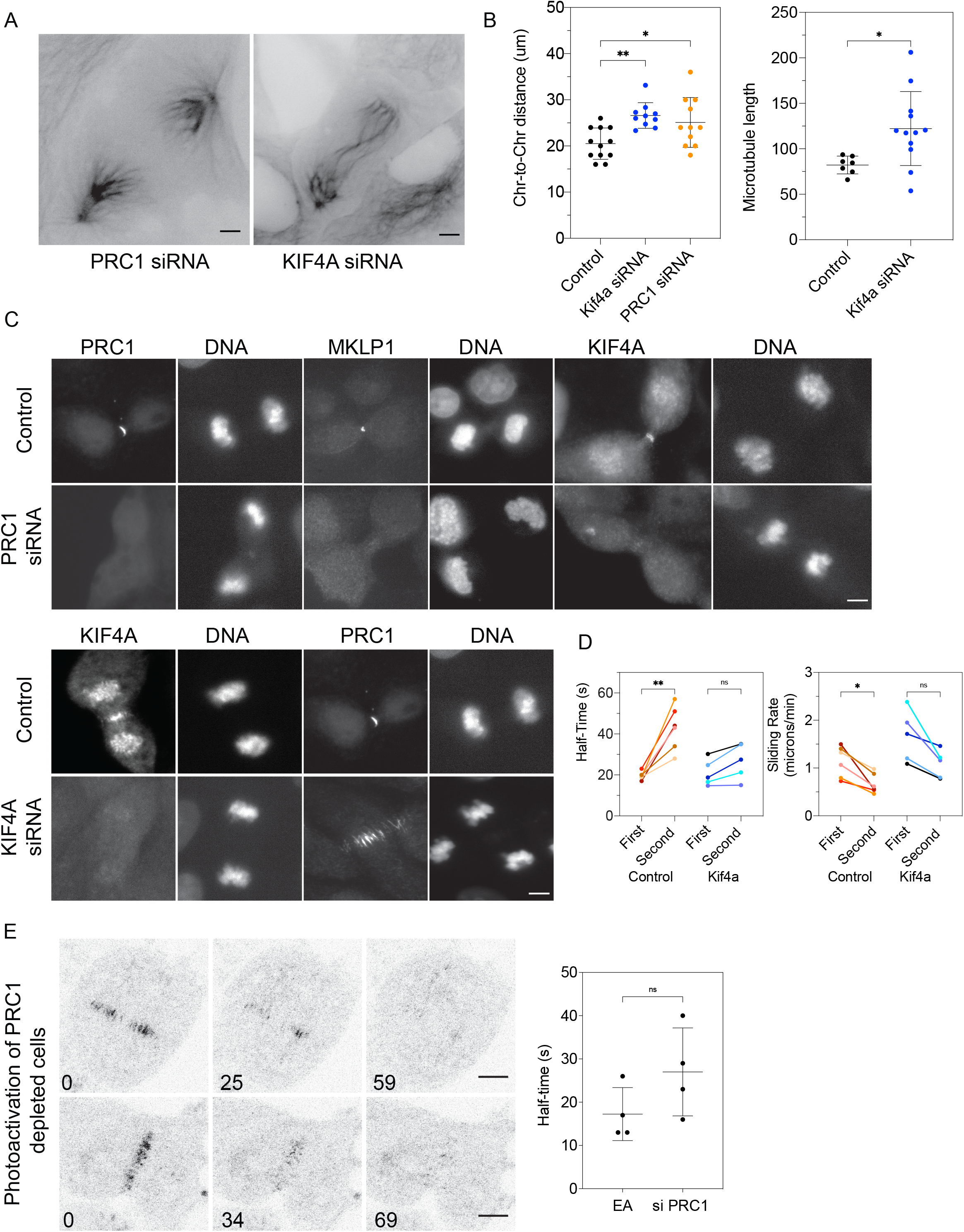
Phenotype of cells depleted of PRC1 or Kif4a. (A) Distribution of midzone microtubules in LLC-Pk1 cells expressing GFP-*α* tubulin depleted of PRC1 (left) or Kif4a (right). Bar = 5 µm. (B) Distance between segregated chromosomes for control, Kif4a depleted and PRC1 depleted cells measured after photoactivation (left panel). Length of midzone microtubules from Z-slices of EGFP-tubulin expressing cells imaged by spinning disc confocal; individual microtubules, or microtubule bundles were measured for control (n = 7 cells) and Kif4a depleted cells (n = 12 cells) (right panel). (C) Distribution of PRC1, MKLP1 and Kif4a in control cells and cells depleted of PRC1 (top panel). Distribution of Kif4a and PRC1 in control cells and cells depleted of Kif4a (bottom panel). Bar = 5 µm. (D) Double photoactivation experiments on control and Kif4a depleted cells. Dissipation half-time (left) and sliding rate (right); each color is an individual cell. Control cells from Figure 1. (E) Photoactivation of PRC1 depleted cells during anaphase and cytokinesis. Bar = 5 µm; time in seconds; Comparison of dissipation half-time of fast population for control early-anaphase (EA) cells and cells depleted of PRC1. Statistical analysis (B (left)) ANOVA with Dunnett’s test; (B (right), F) Mann-Whitney U test; (D) ANOVA with Holm-Šidák test; (E) Paired ANOVA with Holm-Šidák test. p-values: ns = P > 0.12; * = P < 0.05; ** = p < 0.0021; *** = P < 0.0002; **** = P < 0.0001***.

